# Human INCL fibroblasts display abnormal mitochondrial and lysosomal networks and heightened susceptibility to ROS-induced cell death

**DOI:** 10.1101/2020.09.14.295881

**Authors:** Bailey Balouch, Halle Nagorsky, Truc Pham, Thai LaGraff, Quynh Chu-LaGraff

**Affiliations:** Neuroscience Program, Union College, Schenectady, New York, 12309, USA; Department of Biology, Union College, Schenectady, New York, 12309, USA; Drexel University College of Medicine, 2900 W. Queen Lane, Philadelphia, PA 19129, USA; Department of Neurology, Harvard Medical School, Boston Children’s Hospital, Boston, MA 02115, USA; Department of Genetics, Harvard Medical School Boston, MA, 02115, USA

## Abstract

Infantile Neuronal Ceroid Lipofuscinosis (INCL) is a pediatric neurodegenerative disorder characterized by progressive retinal and central nervous system deterioration during infancy. This lysosomal storage disorder results from a deficiency in the Palmitoyl Protein Thioesterase 1 (PPT1) enzyme - a lysosomal hydrolase which cleaves fatty acid chains such as palmitate from lipid-modified proteins. In the absence of PPT1 activity, these proteins fail to be degraded, leading to the accumulation of autofluorescence storage material in the lysosome. The underlying molecular mechanisms leading to INCL pathology remain poorly understood. A role for oxidative stress has been postulated, yet little evidence has been reported to support this possibility. Here we present a comprehensive cellular characterization of human PPT1-deficient fibroblast cells harboring Met1Ile and Tyr247His compound heterozygous mutations. We detected autofluorescence storage material and observed distinct organellar abnormalities of the lysosomal and mitochondrial structures, which supported previous postulations about the role of ER, mitochondria and oxidative stress in INCL. An increase in the number of lysosomal structures was found in INCL patient fibroblasts, which suggested an upregulation of lysosomal biogenesis, and an association with endoplasmic reticulum stress response. The mitochondrial network also displayed abnormal spherical punctate morphology instead of normal elongated tubules with extensive branching, supporting the involvement of mitochondrial and oxidative stress in INCL cell death. Autofluorescence accumulation and lysosomal pathologies can be mitigated in the presence of conditioned wild type media suggesting that a partial restoration via passive introduction of the enzyme into the cellular environment may be possible. We also demonstrated, for the first time, that human INCL fibroblasts have a heightened susceptibility to exogenous reactive oxygen species (ROS)-induced cell death, which suggested an elevated basal level of endogenous ROS in the mutant cell. Collectively, these findings support the role of intracellular organellar networks in INCL pathology, possibly due to oxidative stress.

## Introduction

Neuronal Ceroid Lipofuscinoses (NCL), commonly known as Batten Disease, is presently a group of 14 inherited fatal neurological disorders. Collectively, NCLs affect 1 in 100,000 live-births worldwide, and as many as 1 in 12,500 in countries of Anglo-Saxon descent [1, 2]. Although NCLs are of varying underlying genetic causes, ages of onset and severity, the group shares many similar clinical presentations, most notably the progressive deterioration of the visual and central nervous system, and the accumulation of unwanted autofluorescence storage materials in the lysosomes. The infantile form, INCL, typically presents during infancy at 6-12 months of age with widespread progressive retinal and central nervous system (CNS) degeneration; this leads to the rapid and severe deterioration in cognitive function, vision, motor coordination, and seizures [2-6]. Lifespan is reduced to 8-11 years [3], or as short as 6 years in the most severe cases [7]. While the disease is typically managed with medications to diminish symptom severity, there are currently no curative treatment options or medications that effectively delay disease progression [6].

INCL is an autosomal recessive disease caused by loss of function mutations in the CLN1 gene, residing on chromosome 1p32, which encodes for the lysosomal enzyme Palmitoyl Protein Thioesterase 1 (PPT1) [8]. PPT1 is a hydrolase enzyme responsible for the cleavage of a thioester bond linking long-chain fatty acids to modified cysteine residues in palmitoylated proteins [9-13]. Palmitate and other lipids are covalently coupled to proteins via a thioester linkage with cysteine residues, both of which are necessary for trafficking and membrane anchorage. Cleavage of the lipid from the protein is necessary for degradation [14-18]. In the absence of PPT1 enzyme cleavage activity, degradation of these lipid-modified proteins is deficient, and fatty acid thioesters accumulate in the lysosomes as autofluorescence ceroid or lipofuscin storage materials [4, 9, 13, 16, 19, 20]. The accumulation of ceroid or lipofuscin in lysosomes is characteristic of all subtypes of Batten Disease [21] and is heterogeneous in composition, consisting of proteins, proteolipids and metals [19, 20]. Specifically in INCL neurons, these lipid-protein aggregates appear in the form of granular osmiophilic deposits (GRODs) and are curvilinear, fingerprint, or rectilinear shaped [13, 21, 22] as detected by electron microscopy studies [9, 20, 23]. GRODs have been identified in neurons as well as non-neuronal cell types including lymphocytes [23, 24], fibroblasts [23, 25, 26], and brown adipose tissues [12].

The underlying pathology of INCL and how PPT1 enzyme deficiency leads to neuronal cell death remains relatively not well understood [17]. Oxidative stress and related damage is a common pathological feature of numerous neurodegenerative disorders [27, 28]. Studies using human INCL brains and PPT1 knock-out mice revealed that the loss of PPT1 leads to caspase activated pathway of apoptosis in neurons, presumably due to ER-induced stress responses [10, 16]. Excess storage material from the lysosome may be trafficked back to the ER, activating the unfolded protein response (UPR), causing ER stress [10, 17, 19]. Reactive oxygen species (ROS) are released from the ER in response to stress, triggering mitochondrial-mediated apoptosis, and contributing to neurodegeneration [17, 19, 29]. Neurons exhibit elevated energetic needs and thus depend heavily on oxidative metabolism and produce higher levels of ROS than other cell types, increasing their susceptibility to oxidative stress [29, 30]. Additionally, while PPT1 is localized to lysosomes in all cell types; in neurons, it is also present in synaptic vesicles facilitating the recycling of synaptic vesicles after the release of neurotransmitters. PPT1 deficiency in neurons causes reduced availability of synaptic vesicles at axon terminals, possibly contributing to the progressive neurodegeneration observed in INCL [4, 19].

In this study, using a PPT1-deficient fibroblast cell line derived from a male INCL donor harboring Met1Ile and Tyr247His compound heterozygous mutations, we investigated the link between ROS-induced ER and mitochondrial dysfunction with INCL pathogenesis. Our results indicated that INCL patient fibroblasts exhibited a higher level of autofluorescence storage materials and increased LAMP1 signal. INCL patient cells display organellar pathology, specifically disrupted lysosomal and mitochondrial networks, an increase in LAMP1-positive vacuolation, and a heightened susceptibility to ROS. These results suggested that oxidation damage due to ER and mitochondrial dysfunction contributes to neuronal cell death in INCL.

Using a conditioned media paradigm, we determined whether the presence of normal PPT1 enzyme in culture media would reduce autofluorescence accumulation and organellar disruption in INCL patient cells, thus lessening cellular pathologies. Previous research has used conditioned media to investigate N-acetylgalactosamine-6-sulfatase lysosomal enzyme deficiency in Mucopolysaccharidosis IVA, a lysosomal storage disorder [31]. Similar to N-acetylgalactosamine-6-sulfatase, PPT1 is secreted into the extracellular space to be taken up by neighboring cells, through the mannose 6-phosphate mediated pathway [4]. These properties allow both enzymes to be strong candidates for *in vitro* enzyme replacement through conditioned media supplementation. Our hypothesis is that wild-type conditioned media would attenuate abnormal phenotypes in INCL patient cells due to the presence of soluble functional PPT1 enzyme in the media. Conversely, patient conditioned media (i.e. media obtained from patient cell line cultures), added to wild-type cultures, may provoke an abnormal phenotype due to the presence of secreted toxic factors. Our results supported this hypothesis: patient cells exposed to wild type conditioned media exhibited reduced levels of autofluorescence and LAMP1 signal *in vitro* as compared to comparable patient cells. However, although the disease pathology lessened, patient cells grown in the presence of normal PPT1 enzyme could not be completely restored to wild type cellular level, most likely due to the loss of continuous endogenous PPT1 function.

## Materials and Methods

### Fibroblast Cell Lines and Conditioned Media Protocol

A PPT1-deficient human fibroblast cell line, GM20389, was derived from a nineteen-year old INCL male harboring Met1Ile and Tyr247His compound heterozygous mutations (Coriell Institute, NJ, USA). A human dermal fibroblast cell line, GM05659, was obtained from a healthy donor as a control (Coriell Institute, NJ, USA). Additionally, two established and transformed cell lines, the human foreskin fibroblasts line HFF and human lung fibroblasts line MRC-5, were used as additional controls (ATCC, Inc. VA, USA). HFF and MRC-5 cells were cultured in either DMEM or EMEM supplemented with 4.5 g/L glucose and sodium pyruvate (Corning, Inc. Virginia, USA), 10% Fetal Bovine Serum (FBS) (Atlanta Biologicals, Georgia, USA) 1% penicillin-streptomycin (J R Scientific, Inc, California, USA) and 1% of 200mM L-Glutamine (Life Technologies, California, USA). Cultures were incubated at 37**°** C and 5% CO2.

Cultures of two human fibroblast lines, wild type GM05659 (WT) and INCL patient GM20389 (PT), were used to produce the WT- and PT-conditioned media. Conditioned media was obtained by saving media from either WT or PT two-three days old cultures. Cells were maintained at confluency in T25 flasks for six weeks in order to prevent proliferation and allow for better modeling of the post-mitotic state of neurons. Media was replaced every 2-3 days with 50% fresh media and 50% of either WT- or PT-conditioned media. Four conditioned groups were created - group 1: WT culture receiving 50% WT conditioned media (WT+WT), group 2: WT culture receiving 50% PT conditioned media (WT+PT), group 3: PT culture receiving 50% WT media (PT+WT), and group 4: PT culture receiving 50% PT conditioned media (PT+PT).

### Fixation and Antibody Staining

Early passage cells (passage 2 through 5) were counted using the Scepter automated cell counter (MilliporeSigma, Massachusetts, USA) and cells were plated onto coverslips onto a 12-well plate, at a density of 4 x 10^4^ cells per well, 48 hours prior to fixation for maximum adherence. Cells were then fixed in 4% formaldehyde in Phosphate Buffered Saline (PBS) for 15 minutes at room temperature, followed by three washes with PBS, followed by permeabilization with PBT (PBS with 0.1% Tween-20) and three more PBS washes. Cells were stained with either single or a combination of the following: phalloidin conjugated to Alexa-594 [1:1250] (Thermo Scientific, Germany), phalloidin conjugated to Alexa-488 [1:1000] (Thermo Scientific, Germany), MitoTracker [600nM] (Molecular Probes-Invitrogen, Oregon, USA), or DAPI [1 µg/mL] (MilliporeSigma, Massachusetts, USA) intracellular markers. The following primary antibodies were used: Monoclonal anti-LAMP1[1:200] (antibody #H4A3-s Developmental Studies Hybridoma Bank, Iowa, USA), polyclonal anti-cathepsin D [1:200] (antibody #bs-1615R Bioss Antibodies, Massachusetts, USA), polyclonal anti-vimentin [1:200] (antibody #bs-0756R Bioss Antibodies, Massachusetts, USA), and monoclonal anti-beta-tubulin [1:200] (antibody #86298T Cell Signaling, Massachusetts, USA). Secondary antibodies used were Alexa 488-conjugated goat anti-mouse, Alexa 594-conjugated rabbit anti-mouse, Alexa 488 or Alexa 594-conjugated goat anti-rabbit secondary antibody [1:400] (Invitrogen, Inc. Oregon, USA). Cells were incubated in: Phalloidin for 30 minutes, MitoTracker for 120 minutes, DAPI for 5 minutes, primary antibodies for 60 minutes, and secondary antibodies for 45 minutes. All incubations took place at 37**°** C and 5% CO2. Staining was followed by three additional PBS washes, and a final wash in dH_2_O.

### Fluorescence Imaging and Analysis

Following fixation and staining with the appropriate antibodies or intracellular markers, coverslips were mounted onto microscope slides using anti-fade medium (Molecular Probes-Invitrogen, Oregon, USA), and visualized with a Zeiss AX10 Observer A1 inverted microscope, equipped with a SPOT imaging camera and software (Diagnostic Instruments Inc, Michigan, USA). Fluorescence was observed using DAPI (ex 358nm / em 461nm), GFP (ex 488nm / em 530nm), and Texas Red (ex 596nm / em 620nm) filter channels. Cells were viewed with 100X-oil immersion and 40X objectives.

For quantitative fluorescence imaging, all cells were imaged at 100X magnification with oil immersion objective. The imaging parameters were optimized for every staining. Slides free of previous fluorescence exposure were imaged using identical parameters. To account for photobleaching, all slides were imaged sequentially, and fluorescence exposure was timed and limited to less than 45 minutes per slide. Only non-overlapping cells were imaged for quantitative fluorescence analysis, in order to not mistake combined signal for increased signal intensity. Images were analyzed using the freehand selection tool in ImageJ (NIH, Maryland, USA) to trace the perimeter of cells and obtain measurements of mean and maximum intensity. A square representative of the background signal was also measured for each image. The background signal was subtracted from the mean signal intensity of the cell to be used for analysis.

To count vacuolation, the number of vacuoles in HFF and PPT1-deficient cells stained with LAMP1 were counted using a mechanical hand counter. A minimum of 299 cells were analyzed per conditioned group, across three replicates. Only vacuoles with a defined border were counted. Images of LAMP1-positive vacuoles reported [32] were used as reference. For autofluorescence analysis, a minimum of 121 cells were analyzed per conditioned group, across two replicates, and background intensity was subtracted from the mean signal intensity to correct for heightened background signal due to prolonged exposure time.

### MTT Cell Viability and H_2_O_2_ Induced Cell Death Assay

Cell viability was determined by MTT (C, N-diphenyl-N′-4,5-dimethyl thiazol-2-yl tetrazolium bromide) assay (Roche, Switzerland), a standard colorimetric assay which uses the metabolic reduction of the tetrazolium salt to form the colored formazan product [33]. Here, MTT assay was used to measure metabolic activity as an indicator for cell viability [34]. Cells were counted using a hemocytometer and adjusted to a concentration of 1 x 10^5^ cells/ml. 1 x 10^4^ cells (100µl) were plated per well of a 96-well plate. The number of cells plated was determined based on the finding that 1 x 10^4^ cells were confluent in a 96-well plate upon adherence. Because the assay was used as a viability assay rather than a proliferation assay, cells were plated in 1% serum containing DMEM, in order to measure the reduction in cell viability with 10,000 confluent cells as the baseline. To measure cell viability, cells were given a 48-hour incubation period. After the incubation period, 0.5mg/ml of MTT reagent (10 µl) was added to each well, and the plate was incubated for 4 hours to allow the reagent to be reduced. 100 µl of solubilization solution was added to each well, and incubated overnight. Viability was determined by the absorbance at 550nm minus the reference wavelength 690nm minus a plate blank (as per manufacturer’s protocol). Absorbance was measured using a Spectramax M5 plate reader (Molecular Devices, California, USA). The experiment was repeated with a viability measurement at 120 hours post-plating. To determine cell viability after hydrogen peroxide (H2O2) exposure, 1 x 10^4^ cells were plated per well of 96-well plate, and given 24 hours to adhere. Cells were then treated for 24 hours with 0, 25, 50, or 100 µM H2O2 in DMEM supplemented with 10% FBS, 1% Glutamine, and 1% penicillin-streptomycin. The MTT procedure was carried out as described to determine cell viability after 24 hours H2O2 exposure.

### Detection of ROS

Endogenous ROS levels were measured using a ROS-GLO kit (Promega, Wisconsin, USA) according to manufacturer’s protocol. All four conditioned groups were plated at 2 x 10^4^ cells per well in a 96 well plate and left in the incubator for 24 hours to adhere, prior to ROS detection. Luminescence was measured using a Spectramax M5 plate reader (Molecular Devices, California, USA).

### Statistics

All statistical calculations were performed using Excel (Microsoft, Washington, USA) and SPSS Statistics (IBM, New York, USA). Quantitative fluorescent means were normalized to the WT control of each replicate for both autofluorescence and LAMP1. A one-way ANOVA was used to compare relative fluorescence intensity (RFI) of LAMP1 in PT and WT cells prior to the creation of conditioned groups, with a confidence interval of 99%. A one-way ANOVA with *post-hoc* Tukey HSD test was used to compare RFI of LAMP1 and autofluorescence in conditioned groups; differences were considered significant at a 99.9% confidence interval. A two-sample t-test was performed in ROS detection to compare relative luminescence units, and in LAMP1 signal intensity in early passage PT and WT cell lines. H2O2 induced cell death was analyzed using a univariate ANOVA.

## Results

### PPT1-deficient fibroblasts displayed higher autofluorescence and decreased cell viability

The hallmark of INCL is the presence of autofluorescence storage materials in the lysosomes [13, 21]. We first determined whether autofluorescence signal could be detected in PPT1-deficient human fibroblasts. Autofluorescence signal can be detected in both wild type primary human and INCL patient fibroblasts, and in established human fibroblast controls HFF and MRC using the GFP filter. Autofluorescence was very low, barely detectable in wild type primary fibroblast (n= 50), HFF (n = 56) and MRC-5 (n = 57) controls, but was marginally visible in PPT1-deficient fibroblasts (n = 52) (data not shown). We quantify the fluorescence levels in wild type HFF and MRC fibroblasts and PPT1-deficient fibroblasts to assess whether the difference between normal and patient cells were statistically significant. Upon relative fluorescence intensity (RFI) analysis, PPT1-deficient fibroblasts exhibited a 4.5-fold increase in autofluorescence signal compared to controls. The One-Way ANOVA and *post hoc* Tukey HSD analysis indicated that this RFI increase was statistically significant (p < 0.001). In contrast, RFI between wild type control cell lines did not differ significantly (p = 0.996) (Fig 1).

**Fig 1.**
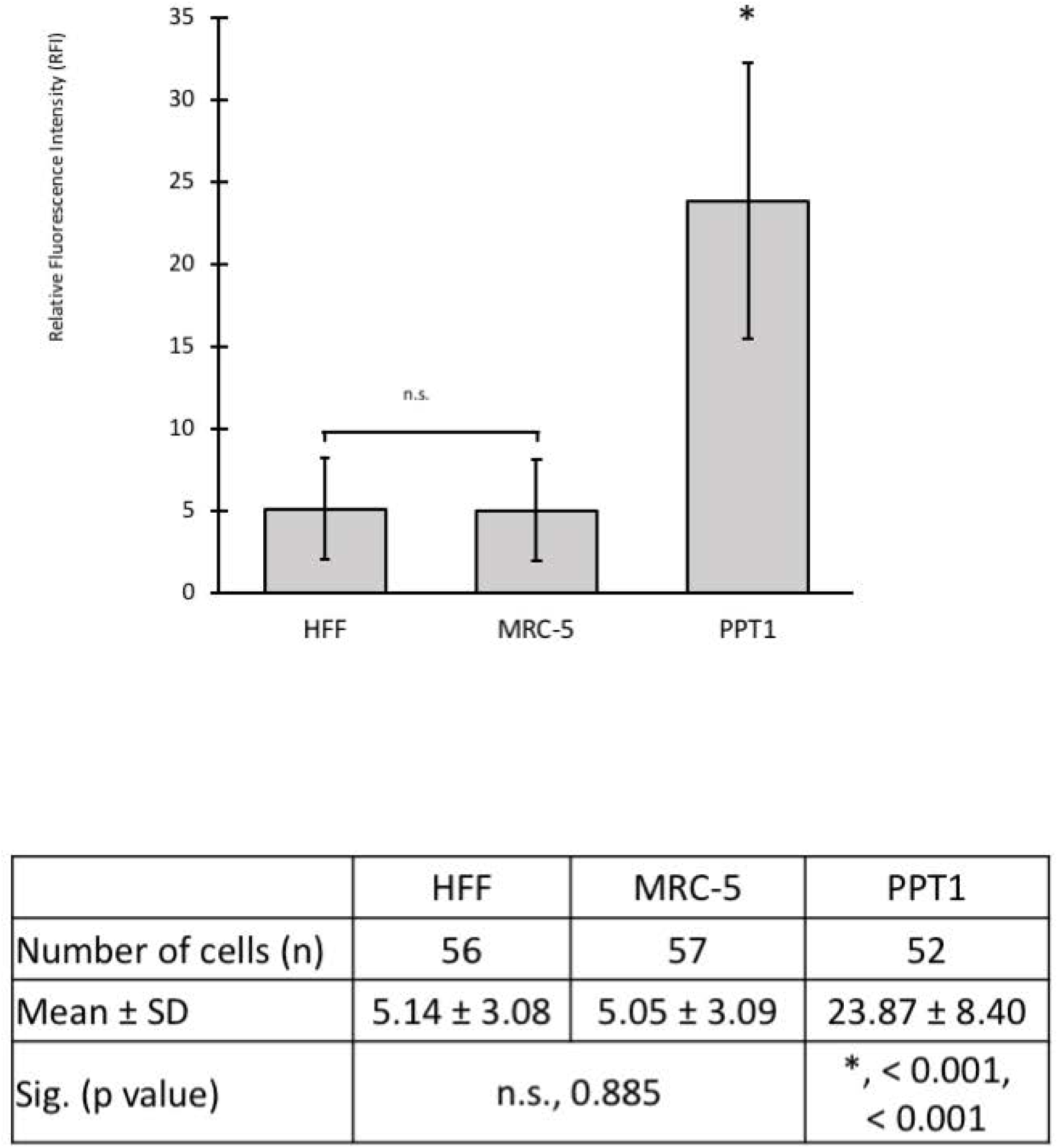
Analysis of autofluorescence in normal and PPT1-deficient fibroblasts. There was a detectable increase in autofluorescence signal in PPT1-deficient fibroblasts compared to HFF and MRC-5 controls. Autofluorescence was increased >4.5-fold in PPT1-deficient fibroblasts compared to controls. Cells were stained with phalloidin-594 at 1:1250 to use as a reference for focusing, and imaged using DAPI (ex 358nm / em 461nm) and GFP (ex 488nm / em 530nm) filters. HFF (n = 56), MRC-5 (n = 57), and PPT1 deficient (n =52) fibroblasts were analyzed by measuring the relative fluorescent intensity using ImageJ. Error bars display +SD. There were no significant differences in relative fluorescence intensity (indicated by “n.s.”) between the HFF and MRC-5 controls (p = 0.885). Fluorescent signal was significantly higher in PPT1 deficient cells as compared to HFF (*, p < 0.001) and MRC-5 (*, p< 0.001) controls.

To investigate the possibility that increased autofluorescence observed in INCL fibroblasts would lead to impaired cell viability, the MTT (C, N-diphenyl-N′-4,5-dimethyl thiazol-2-yl tetrazolium bromide) proliferative assay was used to measure metabolic activity as an indicator for cell viability [34]. PPT1-deficient fibroblast viability was reduced compared to HFF and MRC-5 controls (Fig 2). Specifically, after 48 hours, PPT1-deficient fibroblast viability (n = 7) was reduced significantly compared to that of HFF (n = 8) and MRC-5 (n = 8) controls (p < 0.001). Significant differences were also observed between HFF and MRC-5 controls (p <0.001). This experiment was repeated with an incubation time of 120 hours; the same relative cell viability distribution was observed. PPT1-deficient cell viability (n = 5) was significantly reduced compared to HFF (n = 7) and MRC-5 (n = 7) controls (p < 0.001). Significant differences were also observed between control cell lines (p < 0.001).

**Fig 2.**
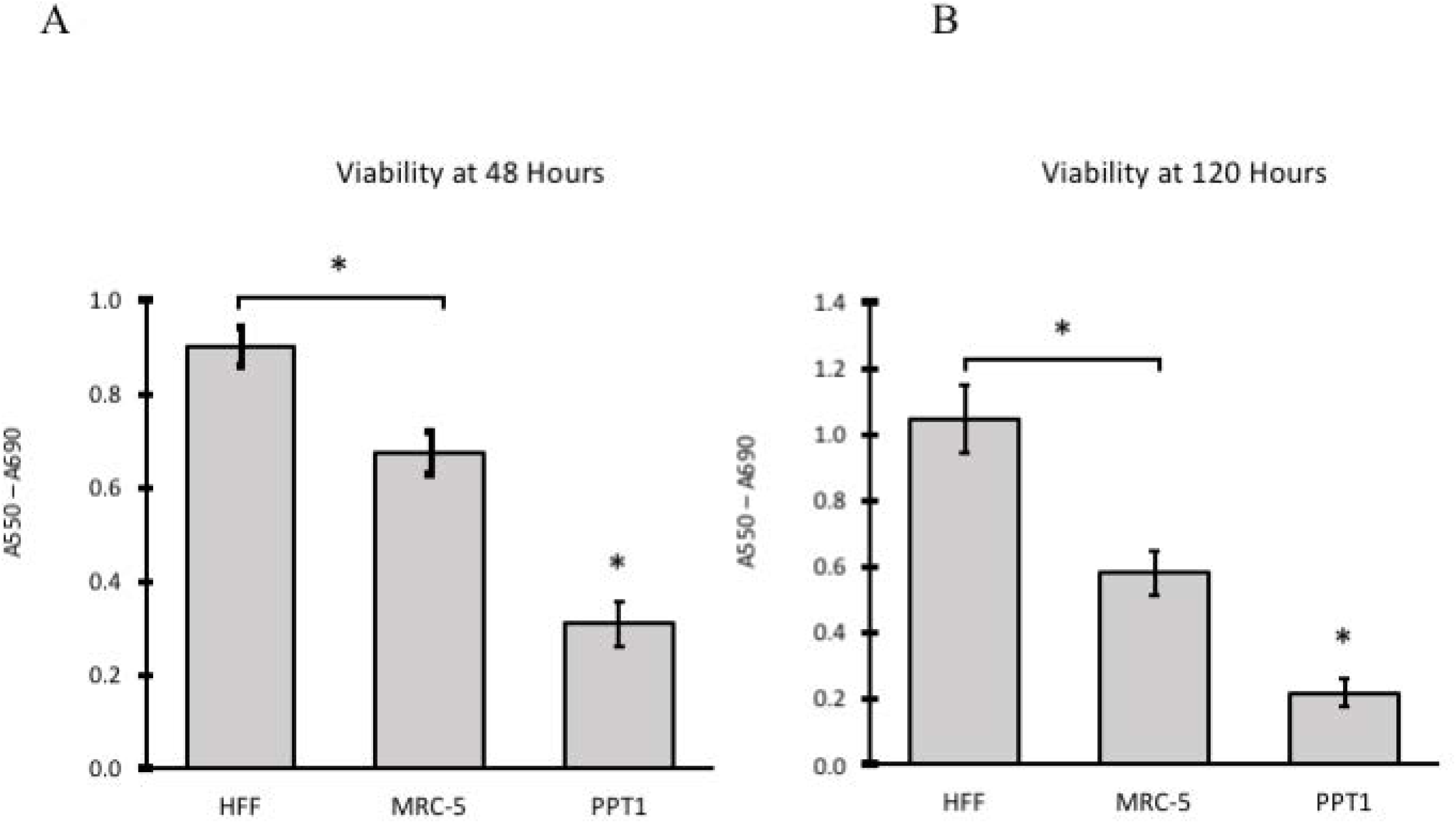
PPT1-deficient fibroblasts display reduced cell viability compared to controls. 1×10^4^ cells from each group were plated in 5 - 8 wells of a 96-well plate in 1% serum DMEM, and viability was determined by MTT assay. (A) Viability was determined 48 hours post-plating. (B) The experiment was repeated with a viability measure at 120 hours post-plating. PPT-deficient cells displayed a similar reduction in relative cell viability regardless of either time points. Significant differences were found between all groups (*, p < 0.001).

### Autofluorescence intensity levels differed significantly between wild type and INCL conditioned groups

Since PPT1-deficient patient fibroblasts exhibit higher autofluorescence and decreased cell viability, we explored the question of whether exposure to functional PPT1 enzyme would have an observable effect on autofluorescence accumulation.

We examined autofluorescence deposit levels in wild type (WT) and INCL patient (PT) fibroblasts exposed to either WT and PT conditioned media (grouped as WT+WT, WT+PT, PT+WT, and PT+PT; see Methods for details). We posited that the autofluorescence pathology may be attenuated in the presence of functional PPT1 enzymes secreted in the media. Imaging and quantitation of autofluorescence deposits was conducted using fluorescence microscopy at excitation 488nm / emission 530nm (GFP) wavelengths (Fig 3).

**Fig 3A.**
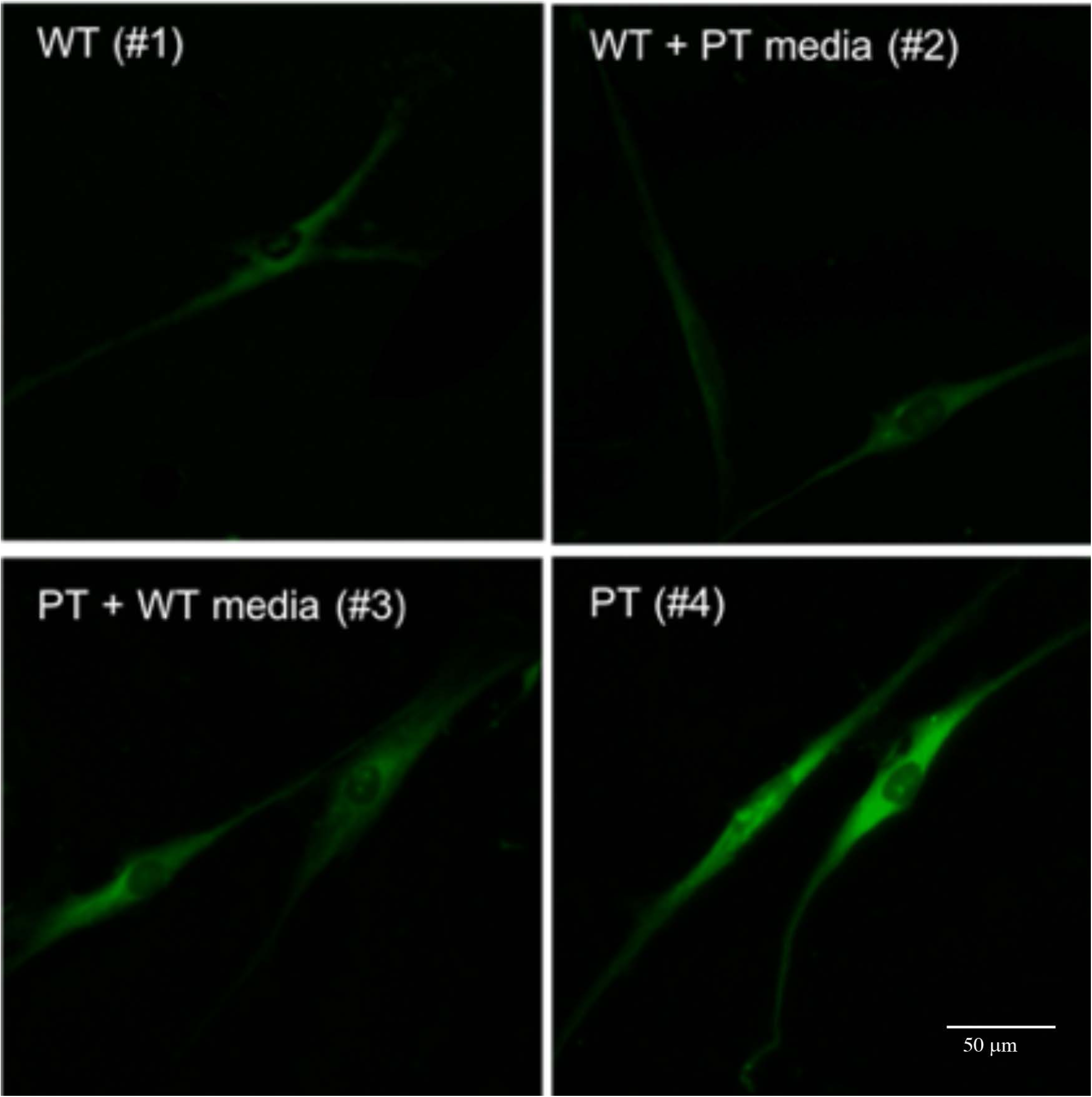
Fluorescence analysis of autofluorescence storage material in four conditioned media groups. Autofluorescence is higher in PPT1-deficient cells grown in either condition 3 or 4 as compared to WT cells grown in either WT or PT-conditioned media (condition 1 or 2). Cells were stained with DAPI at 1:1000 to use as a reference for locating and focusing on cells. Cells were then imaged using the GFP (ex 488nm / em 530nm) filter.

As expected, we observed a three-fold increase in autofluorescence signal intensity (p < 0.001) between the levels of autofluorescence in the WT+WT (group 1) and PT+PT (group 4) treatment groups (Fig 3B). Interestingly, PPT1-deficient fibroblasts exposed to wild type conditioned media displayed significant reduction in autofluorescence as compared to PPT1-deficient fibroblasts incubated in PT-conditioned media (group 3 vs. 4). Signal intensity was decreased to nearly half that of PT+PT cells. Nevertheless, PT+WT cells (group 3) still exhibited 1.63 times greater autofluorescence level than the WT+WT (group 1) indicating that secreted functional enzyme in the media is insufficient to completely restore autofluorescence pathology to normal levels. Additionally, no statistically difference in intensity was observed in WT cells grown in either WT- or PT-conditioned media indicating that endogenous functional PPT1 enzyme were sufficient to overcome potential toxic effects secreted in PT-conditioned media.

**Figure 3B.**
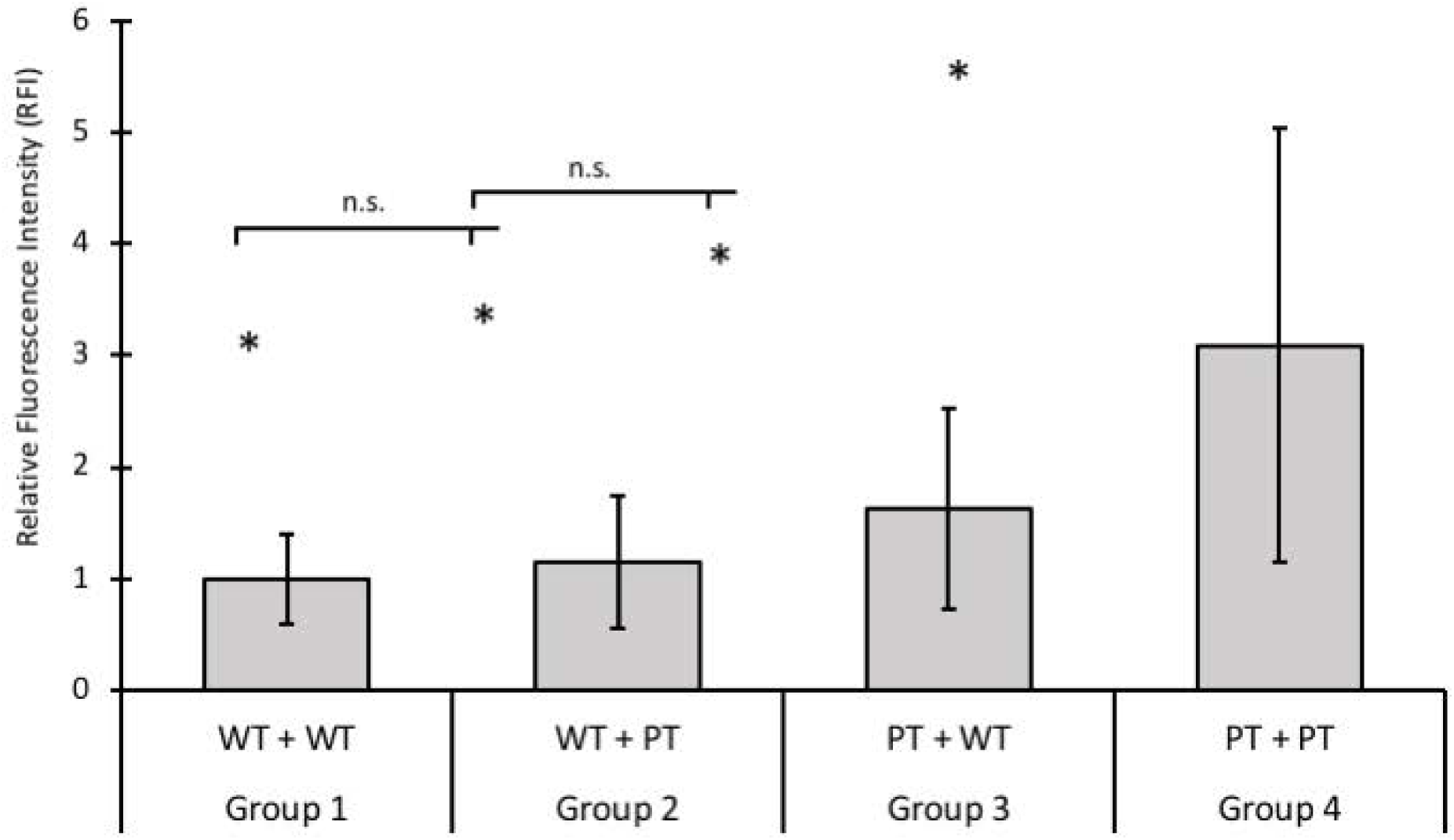
Quantitative analysis of autofluorescence storage material in four conditioned media groups. RFI was measured using ImageJ (n = 2 replicates per group). Significant differences (*, p < 0.001) were found between all conditioned groups, with the exception of group 1 vs 2, and group 2 vs 3, which are labeled n.s (not significant). Error bars indicate +/- SD.

### PPT1-deficient fibroblasts displayed abnormal lysosomal distribution and elevated numbers of lysosomal structures

We next investigated whether autofluorescence storage material was spatially consistent with LAMP1-positive lysosomal structure; and whether increase autofluorescence correlated with abnormal distribution of the lysosomes in PPT1-deficient fibroblasts.

Fluorescence microscopy was performed on primary wild type and PPT1-deficient fibroblasts, and established HFF and MRC-5 fibroblasts revealed that patient fibroblasts exhibited a higher level of lysosomal network as demonstrated by increased LAMP1 staining intensity (Fig 4A). Normal fibroblasts displayed relatively sparse distribution of lysosomes throughout the cell, with a slightly higher concentration of LAMP1-positive lysosomes in the perinuclear region. In contrast, PPT1-deficient fibroblasts exhibited LAMP1-positive lysosomes densely packed throughout the cell body (Fig 4). MFI was compared by one-way ANOVA (p < 0.001) and *post-hoc* Tukey HSD analysis, which showed that LAMP1-positive signal was significantly greater in PPT1-deficient fibroblasts (n = 118) as compared to HFF (n = 114) and MRC-5 (n = 94) controls (p < 0.001) (Fig 4B). Furthermore, a direct examination of LAMP1-positive lysosomal distribution in wild type and PPT1-deficient fibroblasts revealed a detectable difference in fluorescent signal intensity between early passage (P3) PPT1-deficient fibroblasts and wild type fibroblasts (Fig 5A). Analysis showed a statistically significant 1.3-fold increase in LAMP1 signal intensity in PT cells (p < 0.01) (Fig 5B).

**Fig 4.**
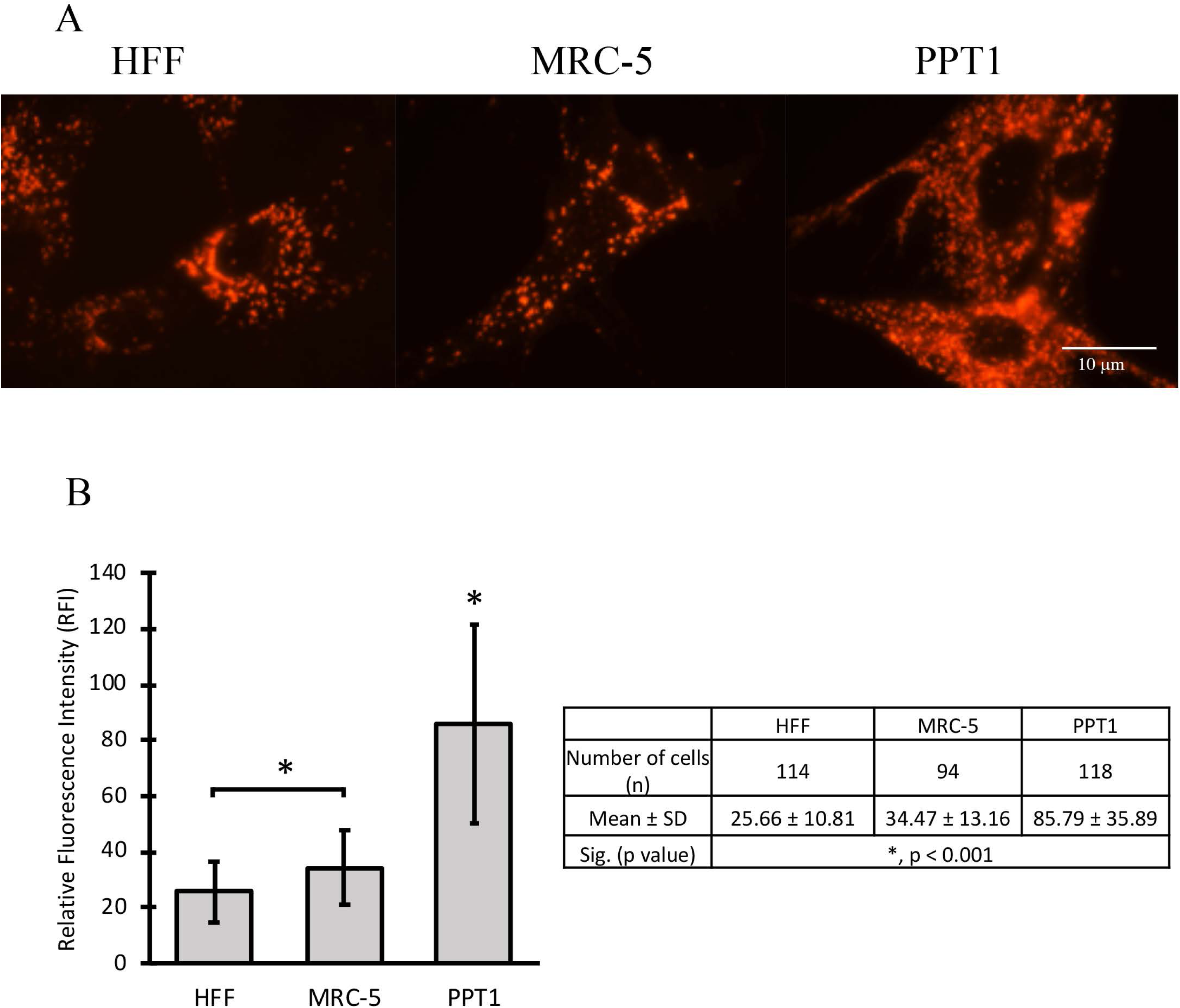
The cytosol of PPT1-deficient fibroblasts is densely packed with lysosomes. (A) HFF, MRC-5 and PPT1 patient cells stained with LAMP1 antibody show the distribution of lysosomes. The density of LAMP1 signal increased in INCL cells as compared to controls. (B) The average LAMP1 signal intensity for the total area occupied by lysosomes was measured using ImageJ (n = 94 - 118 cells per group). Significant differences were found between controls (*, p < 0.001). PPT1 patient cells had significantly higher signal compared to both controls (*, p < 0.001). Error bars display +SD.

**Fig 5A.**
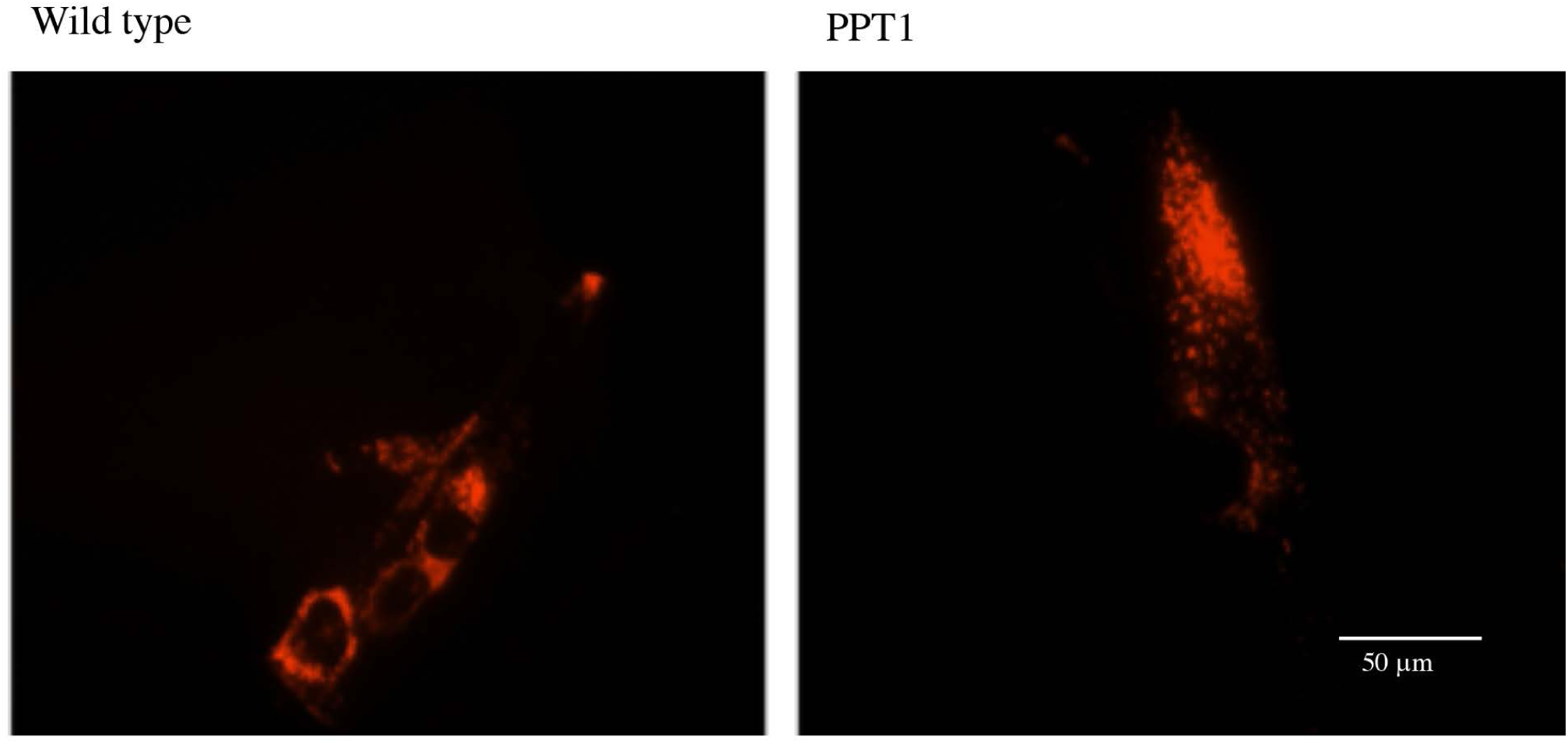
LAMP1 signal in early passage (P3) WT and PPT patient fibroblasts. Cells were stained with LAMP1 antibody, and imaged using a Texas Red (ex 596nm / em 620nm) filter for fluorescence microscopy. LAMP1 signal was greater in PPT patient cells compared to WT (p < 0.01).

**Fig 5B.**
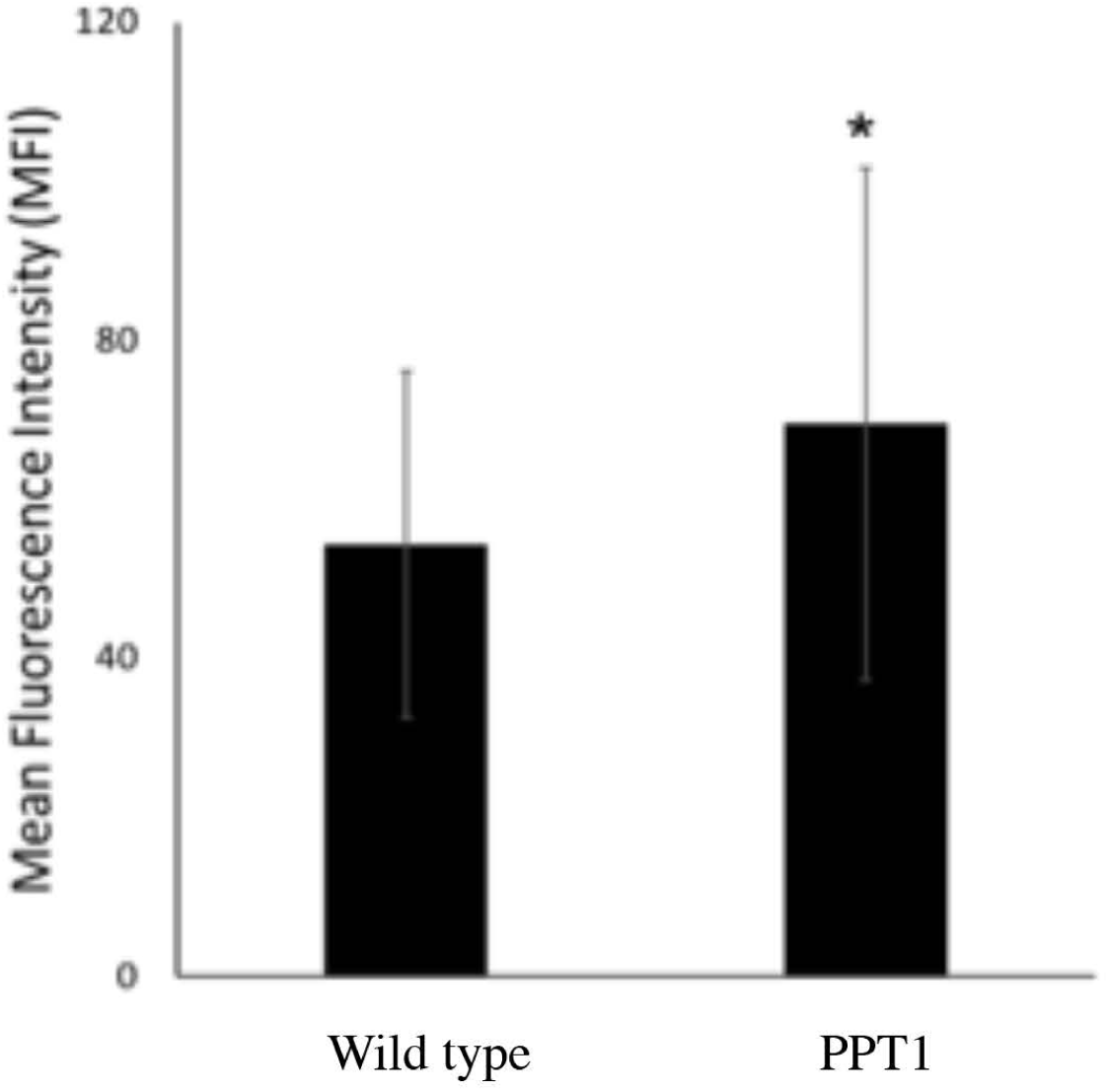
Quantitative analysis of LAMP1 signal in early passage PPT patient and WT fibroblasts. Average LAMP1 signal intensity was measured using ImageJ (n = 40 – 60 cells for both groups). A significant 1.3-fold increase (*, p < 0.01) in mean fluorescence intensity (MFI) was found in PT cells compared to WT. Error bars represent +/- SD.

### Wild type and PPT patient fibroblasts grown in conditioned media exhibited significant differences in the lysosomal distribution

We next assess whether the observed abnormal lysosomal pathology in PPT1-deficient fibroblasts can be lessened in the presence of wild type PPT1 enzyme in conditioned media using LAMP1 antibody staining on all four conditioned groups 1-4 (Fig 6A). LAMP1 fluorescence intensity was statistically significant (p < 0.001) between all conditioned groups (Fig 6B). PT+PT cells (group 4) exhibited a two-fold increase in LAMP1 signal when compared to the WT+WT control (group 1). In contrast, a significant reduction in LAMP1 signal was observed in PT cells grown in WT-conditioned media (group 3) compared to PT+PT cells (group 4), but had a 1.4-fold increase in intensity compared to WT+WT control (group 1). Relative to WT+WT cells, WT cells conditioned with PT media (group 2) were found to have a 1.2-fold increase in LAMP1 signal (Fig 6B).

**Fig 6A.**
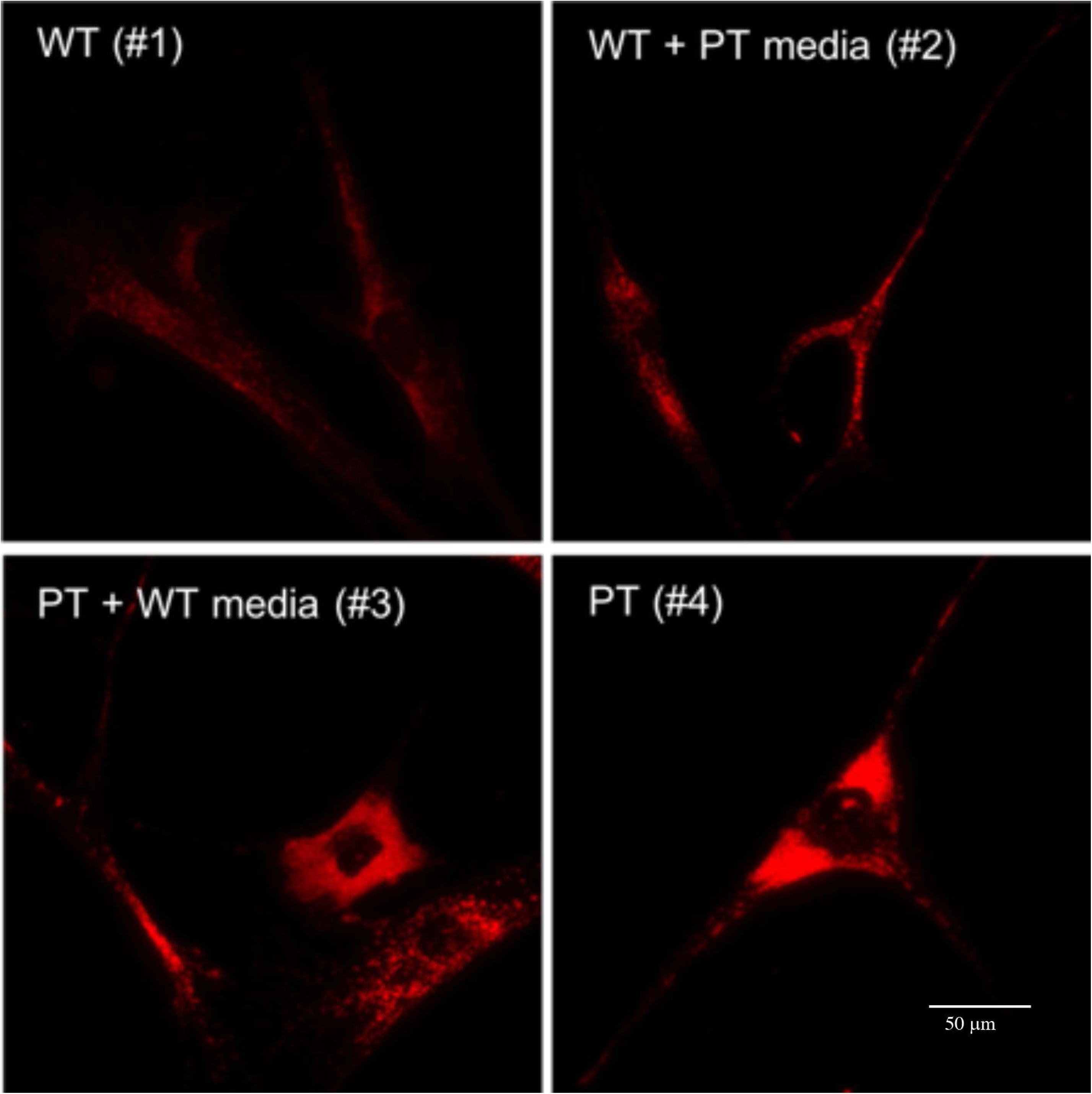
LAMP1 fluorescence signal in four conditioned media groups. Cells probed for LAMP1 were imaged using the Texas Red (ex 596nm / em 620nm) filter for fluorescence microscopy.

**Figure 6B.**
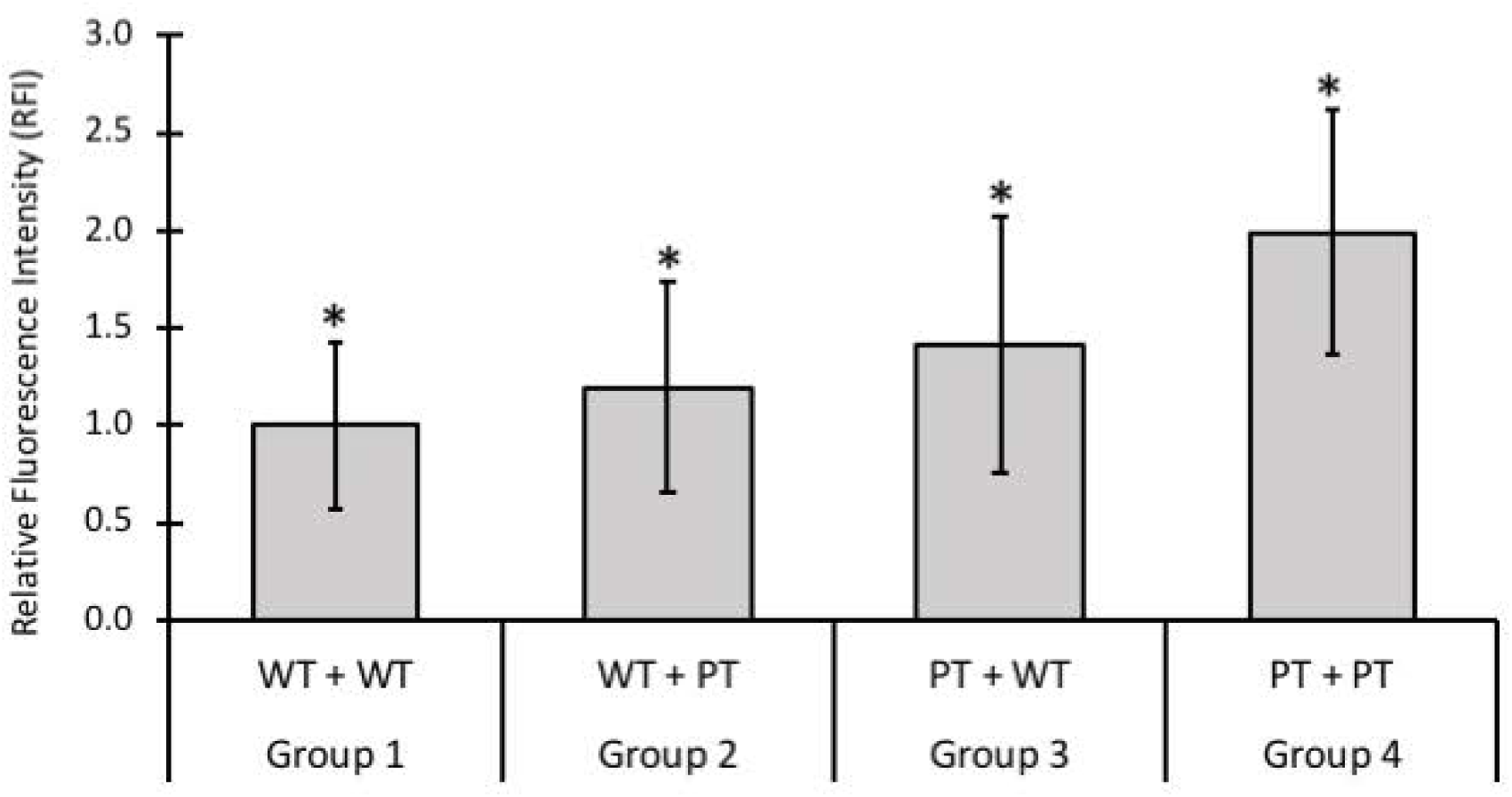
Quantitative analysis of LAMP1 signal intensity in conditioned media groups. Relative fluorescence intensity (RFI) was measured using ImageJ (n = 3 replicates per group). Significant differences (*, p < 0.001) were found between all conditioned groups. Error bars indicate +/- SD.

### PPT1-deficient fibroblasts displayed abnormal mitochondrial network

Lysosomal and mitochondrial dysfunction have previously been associated and implicated in neurodegeneration [27]. Morphological differences have been identified in PPT1-deficient fibroblasts using MitoTracker [26], and evidence of mitochondria-mediated apoptosis has been identified in PPT1 knock-out cells [10]. To follow up on this work, we performed MitoTracker staining to visualize the mitochondrial network in PPT1-deficient fibroblasts and in HFF and MRC-5 controls. Morphologically, mitochondrial tubules could be identified in all control cells observed (Fig 7). MRC-5 cells displayed normal highly branched interconnected tubules. HFF cells had elongated tubules, but also exhibited normal branching (Fig 7 A&B). In contrast, PPT1-deficient cells displayed a substantial decrease in mitochondrial tubule branching, and the mitochondrial network instead consisted predominantly of non-tubular spherical punctate structures (Fig 7C).

**Fig 7.**
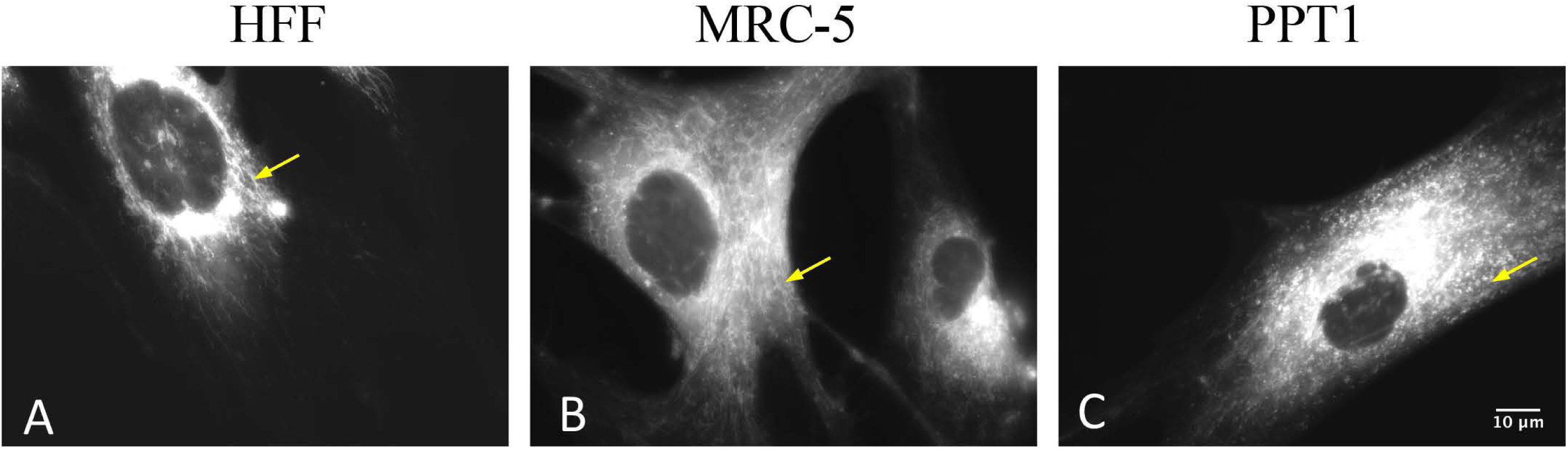
MitoTracker staining shows disruption of the mitochondrial network in PPT1-deficient fibroblasts. Mitochondria staining is observed most heavily in the perinuclear region in all cells. Typical mitochondrial patterning can be seen in controls, where the mitochondria fuse into elongated tubules with extensive branching (arrowhead in A & B). Branching is partially lost in PPT1-deficient cells, and the cytosol is overwhelmed with spherical, punctate structures (arrowhead in C).

### Analysis of lysosomal dysfunction in PPT1-deficient fibroblasts

Because mitochondrial dysfunction leads to impairment of lysosomal activity [27], we sought to determine whether indicators for lysosomal dysfunction could be observed in PPT1-deficient fibroblasts. The enzymatic activity of a lysosomal protease, cathepsin B, has been shown to be decreased due to pharmacologically-induced mitochondrial dysfunction [27]. We analyzed the expression pattern of a closely related lysosomal protease, cathepsin D, in PPT1-deficient fibroblasts because it has been implicated in the initiation of mitochondrial apoptosis [35]. No differences were observed in the relative cathepsin D-positive signal density between PPT1-deficient and HFF control fibroblasts (Fig 8). In both cell lines, cathepsin D-positive signal was observed throughout the body of the cell, and partially within the extending membrane processes. Although overlap between cathepsin D and LAMP1 signal was observed, cathepsin D was not exclusively colocalized to LAMP1. As observed previously, PPT1-deficient fibroblasts displayed a substantial increase in LAMP1 signal as compared to wild type HFF cells (Fig 8). Large LAMP1-positive vacuoles have been shown to form due to ROS produced following mitochondrial damage [27]. We used vacuole formation as an indicator for lysosomal impairment, possibly brought on by mitochondrial dysfunction. The occurrence of large vacuoles formed within LAMP1 stained cells was also increased in PPT1-deficient cells (Fig 8-arrows in bottom right panel). Vacuoles were identified in 43.6% of PPT1-deficient fibroblasts (n = 78), but only 14.1% of HFF (n = 78) controls. Of the cells which had visible vacuoles, the number of vacuoles was also increased to an average of 5 per cell in PPT1-deficient cells versus 2 per cell in the HFF control.

**Fig 8.**
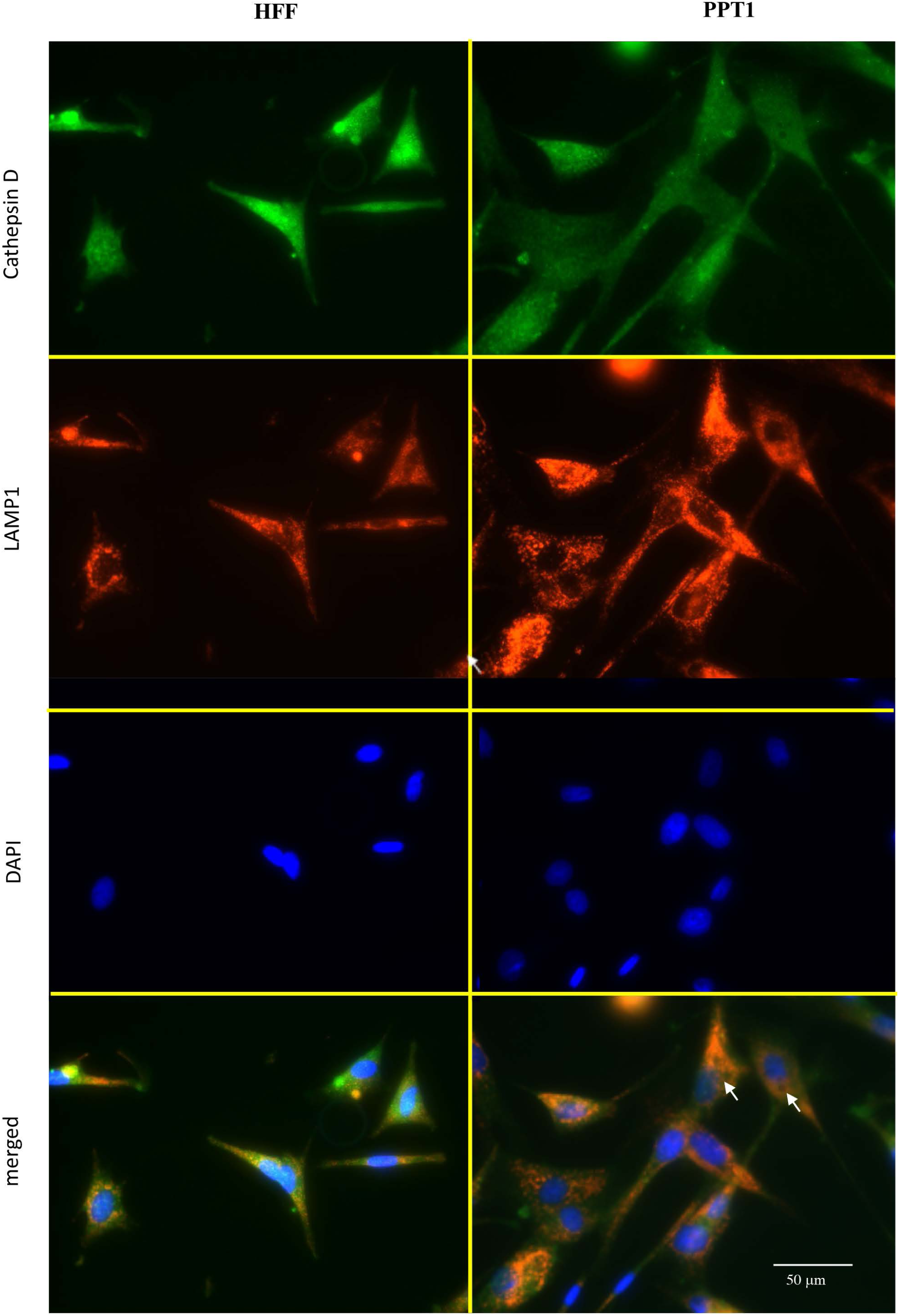
Cathepsin D expression is normal in PPT1-deficient cells. HFF (left column) and PPT1-deficient (right column) cells were stained for cathepsin D (green), LAMP1 (red), and counterstained with DAPI (blue). There were no differences observed in the signal intensity or spatial distribution of cathepsin D in PPT1-deficient cells compared to the HFF control. In both cell lines, cathepsin D was found abundantly throughout the cytosol and was not localized exclusively to the lysosome (indicated by LAMP1-positive structures). White arrows show vacuoles in the cytosol of PPT-deficient cells. Vacuoles were identified in 43.6% of PPT1-deficient fibroblasts (n = 78), but only 14.1% of HFF (n = 78) controls. Of the cells which had visible vacuoles, the number of vacuoles was also increased to an average of 5 per cell in PPT1-deficient cells versus 2 per cell in the HFF control. Images are at 40X magnification.

### PPT1-deficient fibroblasts were more susceptible to H_2_O_2_-induced cell death

The abnormalities found in the mitochondrial network were suggestive of mitochondrial dysfunction [36], which is known to lead to increased ROS production [30]. We then tested whether PPT1-deficient cells would be more susceptible to cell death induced by exogenous ROS, as expected if pre-existing endogenous ROS were present. H2O2 is a ROS with biological significance [37], and treatment with exogenous H2O2 is a well-established assay known to induce apoptosis in a dose-dependent manner [38, 39]. HFF and PPT1-deficient cells were treated with increasing concentrations of 0 to 100 micromolar H2O2 for 24 hours in order to examine susceptibility to oxidative damage by ROS (n = 5 wells per cell line per dose treatment). Control cell viability declined in a dose-dependent manner with increasing H2O2 concentrations. In contrast, PPT1-deficient cell viability was mostly depleted at all tested H2O2 concentration. A univariate ANOVA revealed a significant group x dose effect (p < 0.001); however, group and dose effects individually were not found to be significant (p = 0.071 and 0.054, respectively) (Fig 9A).

**Fig 9A.**
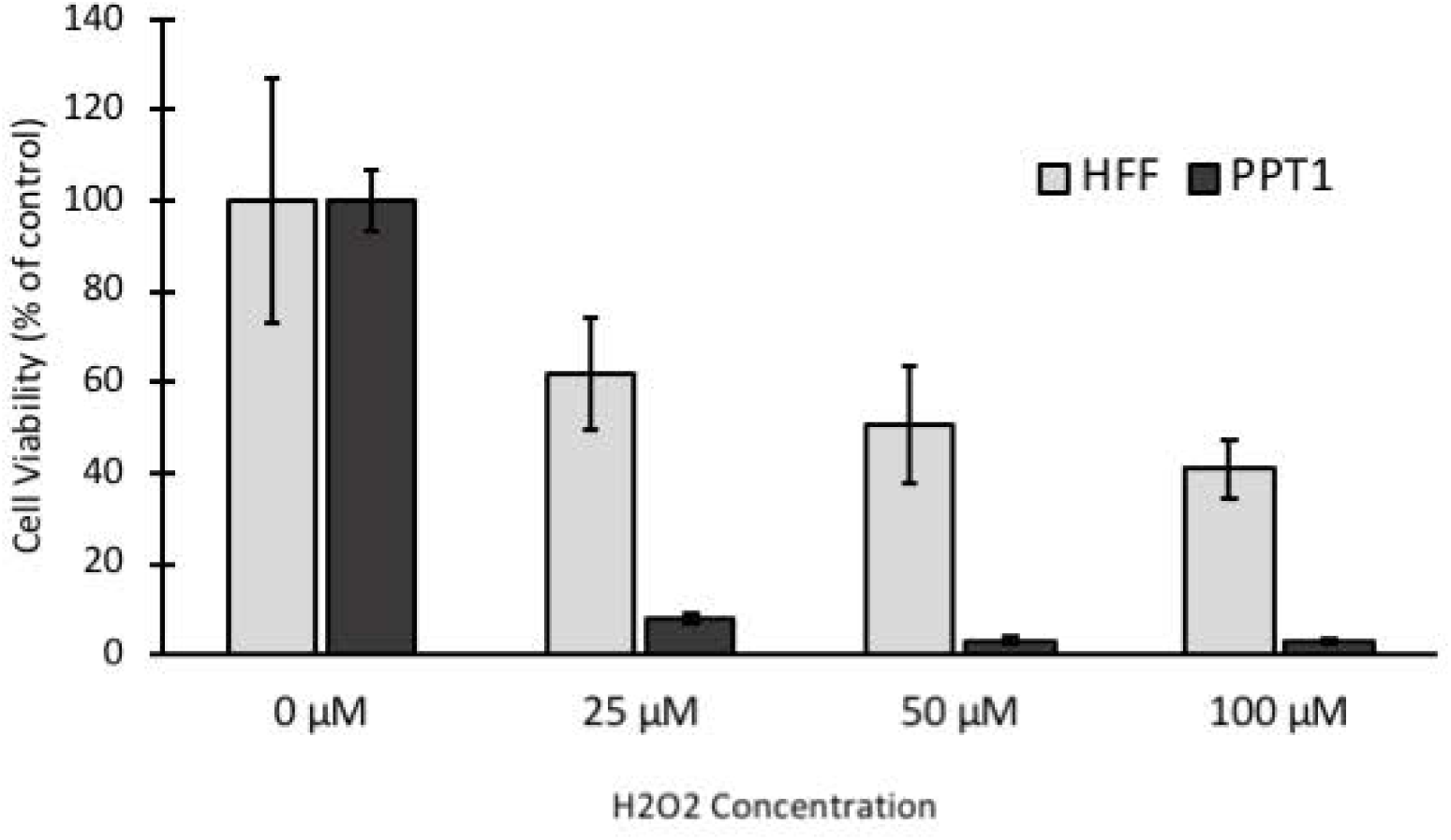
PPT1-deficient fibroblasts are more susceptible to hydrogen peroxide (H_2_O_2_) induced cell death. HFF and PPT1-deficient fibroblasts were treated with 0, 25, 50, or 100 µM H_2_O_2_ for 24 hours. Cell viability was determined by MTT assay. PPT1-deficient cells displayed 8%, 3%, 3% of control viability with the increasing concentrations of 25, 50, and 100 µM H_2_O_2_, respectively. HFF cells displayed 62%, 51%, and 41% of control viability under the same respective conditions. A significant group x dose effect was determined by ANOVA (p < 0.001). Group and dose effects individually were not significant (p = 0.071 and 0.054, respectively). Error bars display +SD.

We also measured endogenous ROS levels in our four conditioned groups 1-4 to ascertain whether the presence of PPT1 in conditioned media had a positive influence on the patient cell’s susceptibility to H2O2 induced cell death. Results indicated while significantly elevated (p < 0.01) relative luminescence units, an indicator for ROS, were detected between both PPT1-deficient groups 3 and 4 as compared to wild type groups 1 and 2, there were not significant difference in the levels of reactive oxygen species whether PPT1 patient cells were grown in wild type (group 3) or PPT1 conditioned media (group 4) (Fig 9B).

**Fig 9B.**
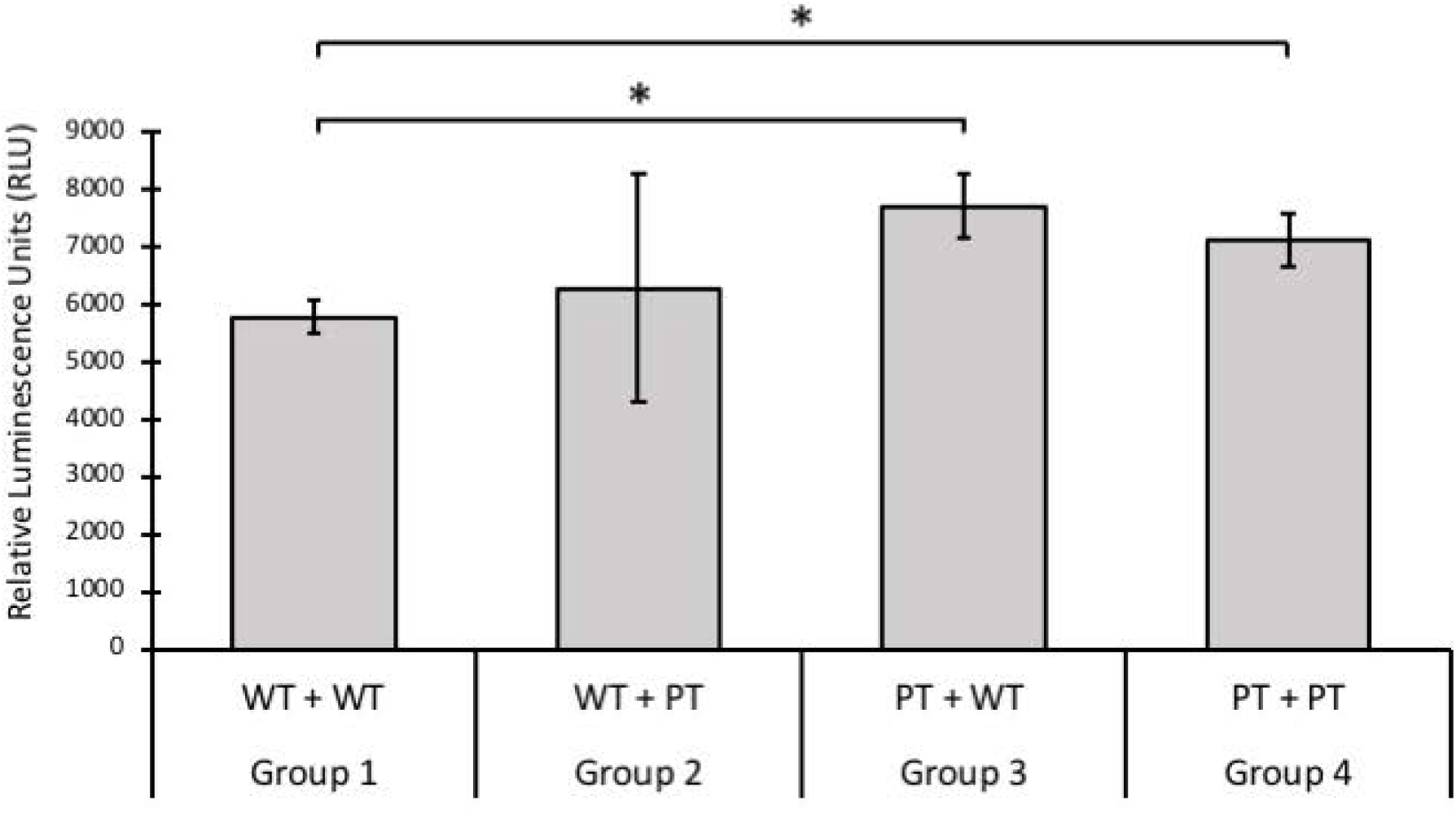
Detection of reactive oxygen species in four conditioned media groups. Intracellular levels of ROS were measured, in relative luminescence units (RLU), using a Spectramax M5 plate reader. A significant increase (*, p < 0.01) in ROS was found in both PT cell groups compared to WT cells. Error bars represent +/- SD.

### The cytoskeleton of PPT1-deficient fibroblasts is morphologically normal

Inhibition of pathways responsible for microtubule assembly has been shown to lead to the accumulation of autofluorescence storage material in the lysosome [21], suggesting an association between components of the cytoskeleton and the lysosome. We examined various components of the cytoskeleton to assess where any distinct morphological abnormalities are observed in PPT1-deficient cells as compared to control cells. Vimentin, a mesenchymal specific intermediate filament, appeared morphologically normal when compared to HFF control cells, as also the microtubule (Supplementary Fig S1). Similarly, the polymerized actin assembly also appeared normal in PPT1-deficient cells (Supplementary Fig S2).

**Supplementary Fig S1.**
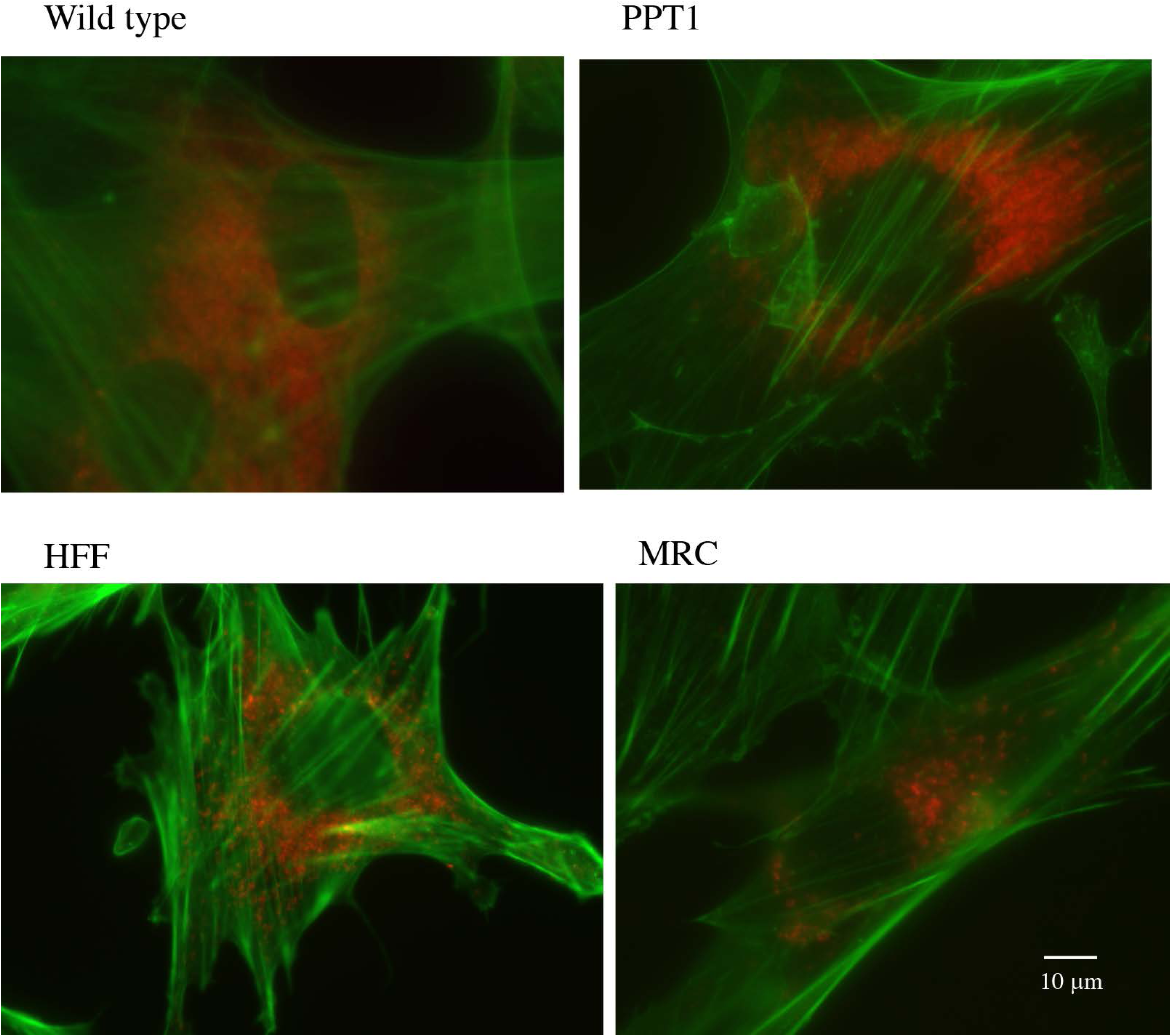
beta-tubulin and vimentin components of the cytoskeleton of PPT1-deficient and normal HFF fibroblasts are indistinguishable. HFF (top row) and PPT1 deficient (bottom row) fibroblasts were probed for vimentin (green), beta-tubulin (red) and counterstained with DAPI (blue). There were no differences observed in the vimentin or beta-tubulin distribution of PPT1-deficient and normal fibroblast cells (merged).

**Supplementary Fig S2.**
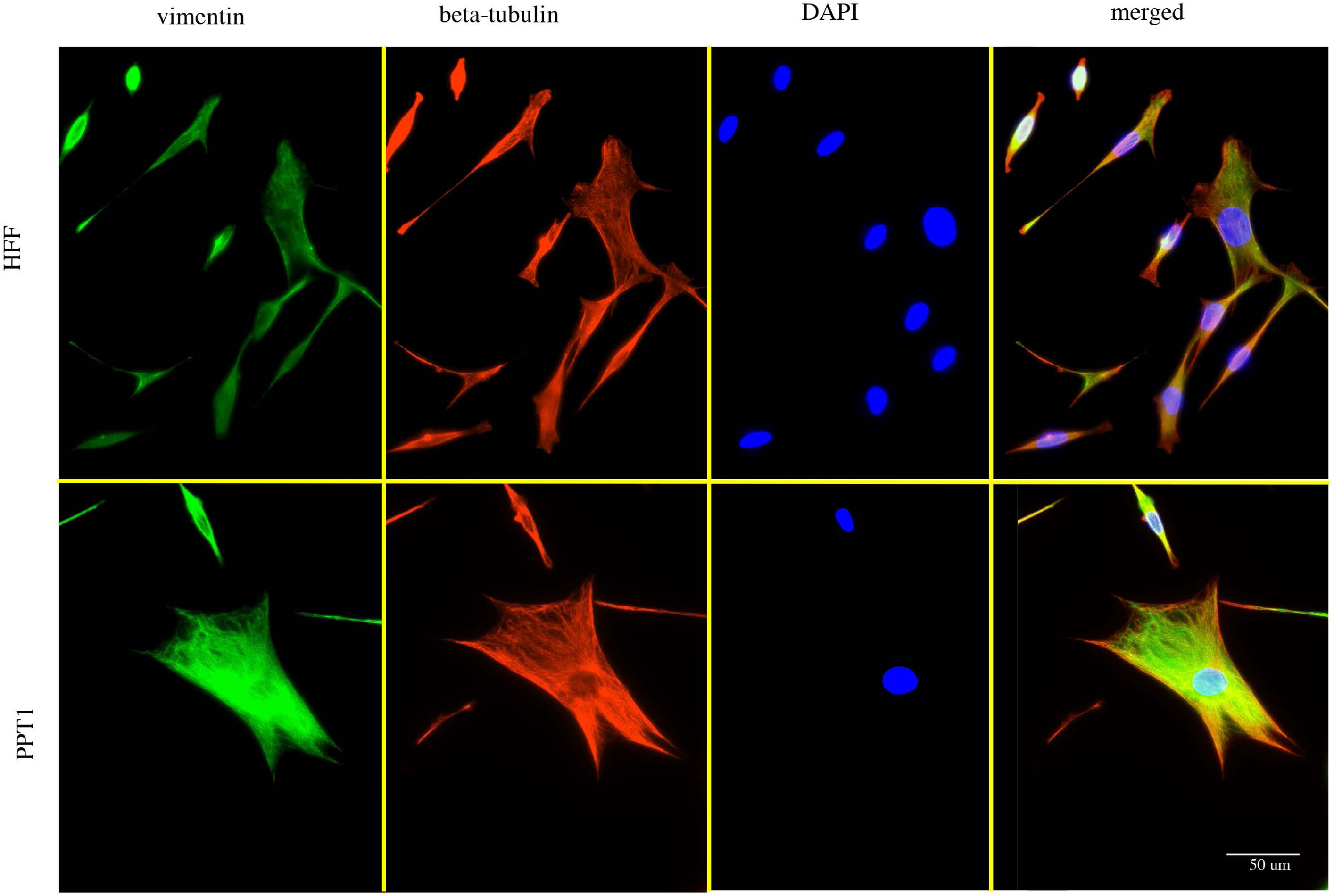
Wild type fibroblasts and PPT deficient fibroblasts exhibit normal actin distribution. Cells were stained with LAMP1 (red) and Phalloidin (green).

## DISCUSSION

Our study describes a detailed characterization of PPT1-deficient fibroblasts derived from a patient with INCL demonstrating that: (a) autofluorescence storage material was present in human fibroblast cells deficient in PPT1 at a level that is higher than wild type; (b) There were organellar pathologies in PPT1-deficient cells, specifically involving the number and distribution of the lysosomal compartments and the mitochondrial network; (c) PPT1-deficient cells had a heightened susceptibility to ROS-induced cell death; (d) There is an increase in LAMP1-positive vacuolation; and (e) The cytoskeleton system, intermediate filaments, microtubules, and actin, were morphologically normal in PPT1-deficient cells indicating that the INCL pathology is discreet and specific to abnormal lysosomal and mitochondrial networks.

Although GRODs are typically detected by electron microscopy [9, 20, 23], we report the detection of autofluorescence storage material in PPT1 deficient fibroblasts using standard fluorescence microscopy - a method similarly used in PPT1-deficient lymphocytes [21], and in brain sections of INCL mice [13]. The presence of autofluorescence storage material also has been reported in TPP1 and CLN3-deficient neural progenitor cells of late-infantile NCL and juvenile NCL [40]. We confirmed the intralysosomal location of autofluorescence storage material in PPT1-deficient patient cells by the co-localization of LAMP1 and autofluorescence signals. This increased autofluorescence accumulation correlates with significantly reduced PPT1-deficient patient cell viability as compared to either fibroblast control cell lines. However, we observed that there are marked differences in cell viability between the MRC-5 and HFF control fibroblasts, most likely due to specificity and robustness of each control fibroblast cell lines. It should also be noted that the PPT1-deficient cells are untransformed and thus are not as robust as either HFF or MRC-5 which may also impact cell viability. It can also be argued that the observed difference reflects levels of metabolic activity rather than direct cell viability. Because the MTT assay uses cell metabolism as an indicator for viability [33], substantial metabolic differences could produce findings which may or may not accurately reflect viability. If this was the case, we believe that decreased metabolic activity reflects a compromised cytosolic state which would eventually lead to lowered cell viability.

Increased lysosomal staining intensity in PPT1-deficient fibroblasts was first reported using the lysosomal marker LysoTracker suggesting an altered pattern of mitochondrial network [26]. Consistent with these findings, we also observed increased staining intensity. Additionally, using LAMP1, we observed a significant increase - three to four folds - in the number of lysosomal structures, as well as dense distribution and localization of the lysosomes beyond the perinuclear region. This indicates that substantially more lysosomal compartments were present in the PPT1-deficient fibroblasts, not just an increase in intensity due to abnormal accumulation and distribution. It has been reported that PPT1 deficiency is closely linked with ER stress and subsequent activation of the ER UPR [10, 18]. PERK is known to play a key role in the activation of the UPR [17], and the transcription factors TFEB and TFE3 have recently been shown to activate lysosome biogenesis in a PERK-dependent manner [41]. The significant increase in lysosomal compartments observed in our study may then represent evidence of increased lysosomal biogenesis due to activation of the UPR, which supports the role for the ER in INCL pathology reported previously [10, 17, 18]. Lysosome biogenesis occurred in a PERK dependent manner which mediates ROS production and activation of mitochondrial-mediated apoptosis in response to ER stress [17].

Our data supports the role of mitochondrial damage in INCL pathology. We observed altered morphology of the mitochondrial network consisting of fragmented mitochondrial tubules and the loss of mitochondrial tubule formation, all of which are indicative of mitochondrial dysfunction. Large spherical mitochondria can arise due to impairment affecting the mitochondrial architecture at the nanoscale [36], and we have observed the spherical punctate mitochondrial morphology in PPT1-deficient patient cells. Since mitochondrial damage and subsequent caspase-9-initiated apoptosis have been implicated in INCL mouse model, and were shown to be ROS-dependent [16], we next sought to investigate whether evidence for ROS existed in PPT1-deficient human fibroblasts. Although ROS has been detected using a PPT1 KO mouse model [10], ROS has not yet been reported in human cells or non-neuronal cell types. Our results indicate that PPT1-deficient human fibroblasts exhibit a heightened susceptibility to cell death induced by exogenous ROS. This is highly suggestive that elevated pre-existing endogenous ROS are already present in PPT1-deficient patient cells.

To determine whether mitochondrial dysfunction could further impair lysosomal function, we assessed the intracellular distribution of cathepsin D and the presence of large vacuole formation from LAMP1-positive cells. Interestingly, cathepsin D was found abundantly throughout both PPT1-deficient and control cells. By qualitative analysis, there were no differences in the spatial distribution or expression of cathepsin D. Previously work has shown that cathepsin D is involved in early stages of the mitochondrial-mediated apoptotic cascade [42]. Cathepsin D-deficiency has also been shown to lead to the accumulation of autofluorescence storage material and progressive cell death, characteristic of the NCLs in general [42, 43]. We find no correlation with cathepsin D density and distribution and mitochondrial dysfunction. Since vacuolization is ROS independent and has no morphological effects on the mitochondrial network [32], the vacuolization observed in PPT1-deficient fibroblasts may represent a direct effect of autofluorescence storage material accumulation on lysosome function rather than a complex interaction with the mitochondria.

Finally, our work indicates that the loss of PPT1 enzymatic activity can be somewhat mitigated with the introduction of wild type PPT1 enzyme in the cytosol. This method was first reported as a potential enzyme replacement therapy for the lysosomal storage disorder Mucopolysaccharidosis IVA [31] and may yet be a similarly viable avenue for INCL. Using this paradigm, we ask whether the media collected from wild type cultures and PPT1-deficient cultures have a positive or negative effect on wild type and PPT1-deficient patient fibroblasts. The introduction of a functional enzyme secreted from wild type conditioned media may restore normal enzyme activity by the observable reduction in autofluorescence storage materials. Alternatively, patient conditioned media when added to wild-type cultures, may provoke an abnormal phenotype due to the presence of secreted toxic factors. Our data indicates that the patient cells benefitted from growing in the presence of wild type conditioned media: there are dramatic reductions in autofluorescence accumulation and LAMP1 positive lysosomes as compared to patient cells grown in their own conditioned media. Although cellular pathology was partially mitigated, restoration was not at the wild type level. Levels of reactive oxygen species were at comparably high levels whether PPT1-deficient fibroblast cells were grown in wild type or PPT1-deficient conditioned media. These results indicate that complete rescue most likely requires constitutive intracellular expression of PPT1 via a gene therapy vector or the direct introduction of the enzyme to the brain or spinal cord. Currently, enzyme replacement therapies for CLN diseases are invasive: intrathecal and intravenous administration of PPT1 in the Ppt1-mouse model spinal cord, and the intracerebroventricularly administration of TPP1 enzyme for the treatment of NCL type 2 [1, 44, 45]. Our study indicates that normal PPT1 enzyme can be internalized by PPT1-deficient cells and be taken up by the lysosomes to repress autofluorescence accumulation and abnormal lysosomal morphology. This paradigm has clinical significance in that a partial cellular recovery may be possible using this passive method.

## Acknowledgments

We would like to thank Union College Undergraduate Funding resources.

